# TASI: A software tool for spatial-temporal quantification of tumor spheroid dynamics

**DOI:** 10.1101/164921

**Authors:** Yue Hou, Jessica Konen, Daniel J Brat, Adam I. Marcus, Lee Cooper

## Abstract

Spheroid cultures derived from explanted cancer specimens are an increasingly utilized resource for studying complex biological processes like tumor cell invasion and metastasis, representing an important bridge between the simplicity and practicality of 2D monolayer cultures and the complexity and realism of *in vivo* animal models. Temporal imaging of spheroids can capture the dynamics of cell behaviors and microenvironments, and when combined with quantitative image analysis methods, enables deep interrogation of biological mechanisms. This paper presents a comprehensive open-source software framework for <u>T</u>emporal <u>A</u>nalysis of <u>S</u>pheroid Imaging (TASI) that allows investigators to objectively characterize spheroid growth and invasion dynamics. TASI performs spatiotemporal segmentation of spheroid cultures, extraction of features describing spheroid morpho-phenotypes, mathematical modeling of spheroid dynamics, and statistical comparisons of experimental conditions. We demonstrate the utility of this tool in an analysis of non-small cell lung cancer spheroids that exhibit variability in metastatic and proliferative behaviors.

## INTRODUCTION

3D spheroid models of cancer have been widely used to investigate mechanisms of invasion and metastasis [1-4] and the impact of drugs on metastatic potential [5-7]. Spheroid cultures help to bridge the gap between simplistic 2D in vitro cultures and complex in vivo mouse models, and have been used to study complex biological processes that are strongly coupled to tissue microenvironments. Cell-cell and cell-matrix interactions in spheroid cultures are more similar to animal models and human disease than 2D in vitro models, yet spheroids can be grown rapidly, are relatively inexpensive and are easier to image than in vivo models. The relative ease in imaging spheroid models makes them especially amenable to investigating temporal processes where dynamic behaviors and interactions can be captured. Metastatic and invasive processes are fundamentally dynamic, and temporal imaging of spheroids can provide important insights into how cancer cells divide [8], invade, and metastasize [1, 5, 9, 10]. For example, measuring growth kinetics of tumor spheroids has been used for anti-cancer drug screening [5, 7]. Co-culturing of multiple cell types in 3D spheroids has also been used to investigate cell-cell interactions in microenvironments [11, 12].

Software for spheroid image analysis has largely focused on static images generated by high throughput screening [4-6, 13-27]. Existing software programs for analyzing spheroid imaging are described in Table 1. Software for measuring spheroid dynamics has received relatively less attention [4, 13, 23-29]. An interactive system for segmenting and measuring spheroid volume and dimensions was developed in [26]. Of the software packages available for spheroid analysis, none provides end-to-end statistical analysis and visualization of imaging measurements. The primary challenges in measuring spheroid dynamics are in accurate delineation of the spheroid boundaries, and the extraction of spatiotemporal features that describe spheroid growth, shape, and motion. Cultures derived from neoplastic cells often exhibit irregular shapes, chaining and branching behaviors, and can be highly dynamic, making automatic delineation difficult [5, 22, 25], and leading investigators to perform manual segmentations that are not objective or repeatable [30, 31]. Similarly, variations in dynamic behavior complicate the extraction of descriptive features. Most spheroid analysis software only measures basic size and shape features, which is insufficient to discriminate different patterns of invasion [32]. Finally, there is a gap between mathematical modeling of behavior following feature extraction, with both capabilities often not available in the same tool [33, 34].

**Table 1.** Summary of software available for spheroid image analysis.

| Software | Category | Analysis & visualization | Availability & language |
| --- | --- | --- | --- |
| **TASI** | Time-lapse | Yes | Open source, Matlab |
| **qVISTA** <sup>[18]</sup> | High-throughput | Yes | Software not provided, Matlab |
| **AMIDA** <sup>[6]</sup>,<br>**AnaSP** <sup>[15, 20]</sup> | High-throughput | No * | Open source,<br>Java <sup>[6]</sup> or Matlab <sup>[15, 20]</sup> |
| **PCaAnalyser** <sup>[21]</sup> | High-throughput | No | Open source, ImageJ |
| **Phaedra** <sup>[14]</sup>,<br>**MetaXpress** <sup>[17, 19]</sup> | High-throughput | No | Commercial |
| **Spheroid Analyzer** <sup>[23]</sup> | Both | No * | Open source, ImageJ |
| **SpheroidSizer** <sup>[26]</sup> | Both | No | Open source, Matlab |
| **Celigo** <sup>[5, 22, 25]</sup>,<br>**Imaris & Velocity** <sup>[24]</sup>,<br>**VTT_Acca** <sup>[4]</sup> | Both | No <sup>[5, 22, 24, 25]</sup>,<br>No * <sup>[4]</sup> | Commercial |

\* Requires other statistical/visualization software, such as R, Excel or additional Matlab functions.

Leader and follower cells were defined in the collective migration process, in which leader cells were migrating at the leading edge of the collective sheet or spheroid, whereas follower cells followed or attached to the leader cells. Leader and follower cells showed distinctive morphology and migration behaviors. For example, leader cells had larger size [29] and were more mesenchymal-like [35]. They formed large lamellipodium and migrated with persistent protrusion [36]. Follower cells were smaller [29] and more epithelial-like [35]. Although leader and follower cell migration behavior were widely studied, most were conducted in 2D models and there were no standard features selected to quantify the morphology differences between them. Furthermore, many researchers still used manual selection to quantify the morphology features of these cells, which limited the number of cells studied and was time-consuming and subjective.

In this paper, we describe TASI, an open-sourced software framework for end-to-end Temporal Analysis of Spheroid Imaging (http://github.com/cooperlab/TASI). This framework allows investigators to automatically segment spheroid images, extract features describing their shape, growth, and invasiveness, and to perform mathematical modeling and statistical analyses of these features to compare treatment and control populations of spheroids. This approach improves the efficiency and objectivity of investigations utilizing spheroid models, and is open-source and extensible by the research community. We demonstrate the utility of this framework with an analysis of lung cancer spheroids and show TASI can discriminate different invasive phenotypes.

## MATERIALS AND METHODS

An overview of the TASI framework is presented in Figure 1 (see supplement Figure S1 for details). Spheroids are first imaged under variable experimental conditions to produce 4D volumes (*x, y, z, time*). Images are then processed using a segmentation algorithm to delineate spheroid boundaries and features are extracted to describe their spatiotemporal characteristics. The temporal evolution of features is described using mathematical models, and statistical tests are performed to compare parameters across conditions, and visualizations are generated. These steps are described in greater detail below.

**Figure 1.**
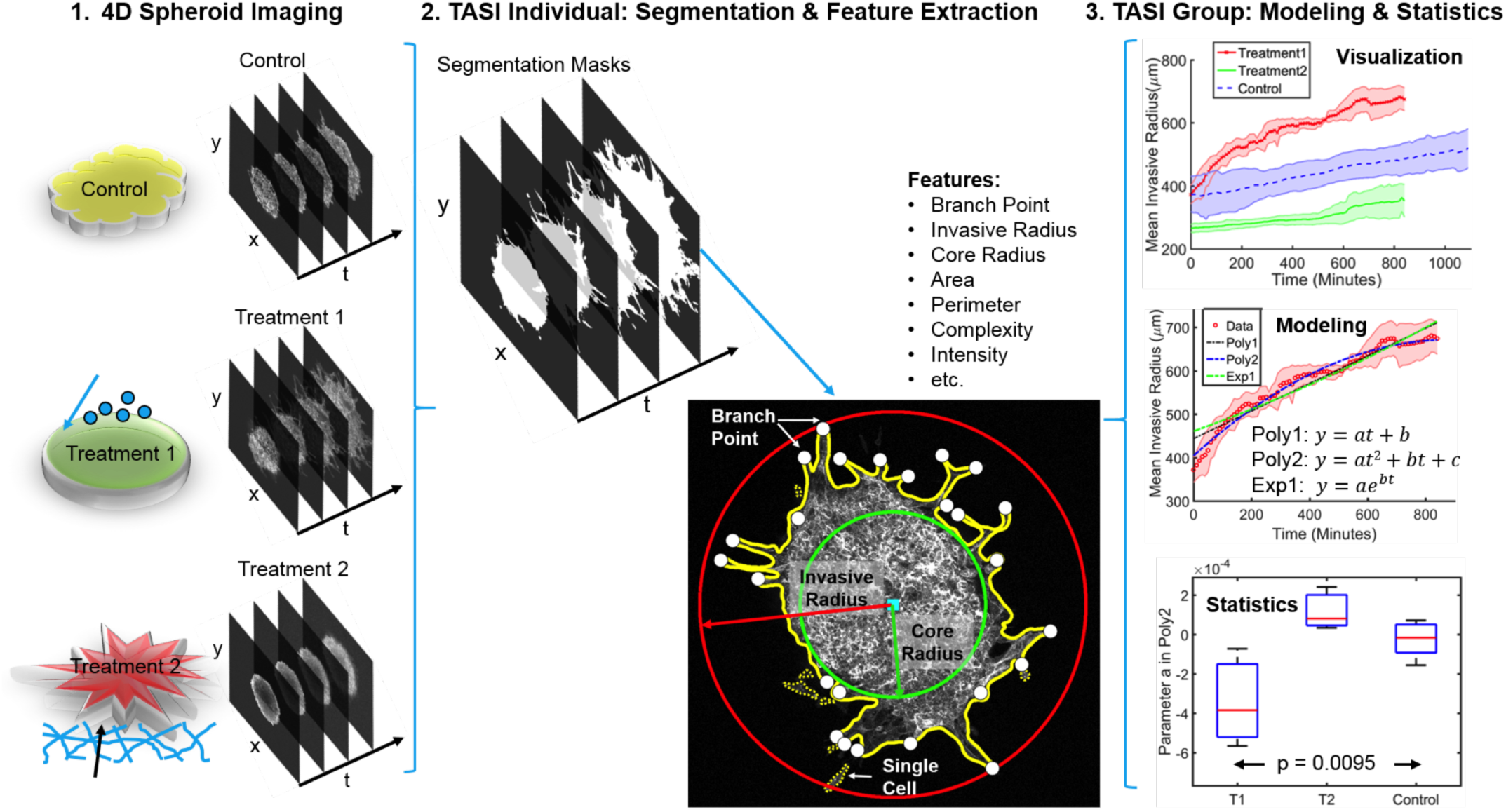
Overview of TASI framework. Step1 shows 4D (x, y, z, t) spheroid dynamic experiment and imaging. Step 2 shows the segmentation and morphology features extracted from the individual spheroid. Solid yellow line is the spheroid boundary. Dashed yellow lines are single cell boundaries. Red circle and red arrow represent the invasive radius. Green circle and green arrow represent the core radius. White dots are branch points and cyan square is centroid of the spheroid. Step 3 show the visualization, modeling and statistical analysis for grouped spheroid dynamics under different treatments.

### Spheroid culture and imaging

Data for validating the TASI framework was generated using the <u>S</u>p<u>A</u>tiotemporal <u>G</u>enomic and cellular <u>A</u>nalysis (SaGA) technique to create spheroid cultures that represent the different invasive and metastatic phenotypes observed in lung cancer [37]. First, parental H1299 non-small cell lung cancer spheroids were embedded in a 3D matrix and visualized using a Leica SP8 confocal microscope. These spheroids contain cells that exhibit a variety of invasive tendencies, forming chain-like growths containing multiple cells that project from the main spheroid body (see example Figure 2.a). The cells at the tips of these chains were isolated using SaGA and cultured to form “leader” cell spheroids (see example Figure 2. a). The cells following the leaders within the chain-like projections were similarly isolated to produce “follower” spheroids (see example Figure 2. a). Each spheroid was imaged at x, y, z planes every 10 minutes for a minimum 14 hours. A total of 6 parental spheroids, 3 leader spheroids and 3 follower spheroids were imaged for the purposes of generating data to validate the ability of our software to quantify cancer invasion and metastasis. Detailed experimental methods for cell culture and imaging are provided in supplementary methods.

**Figure 2.**
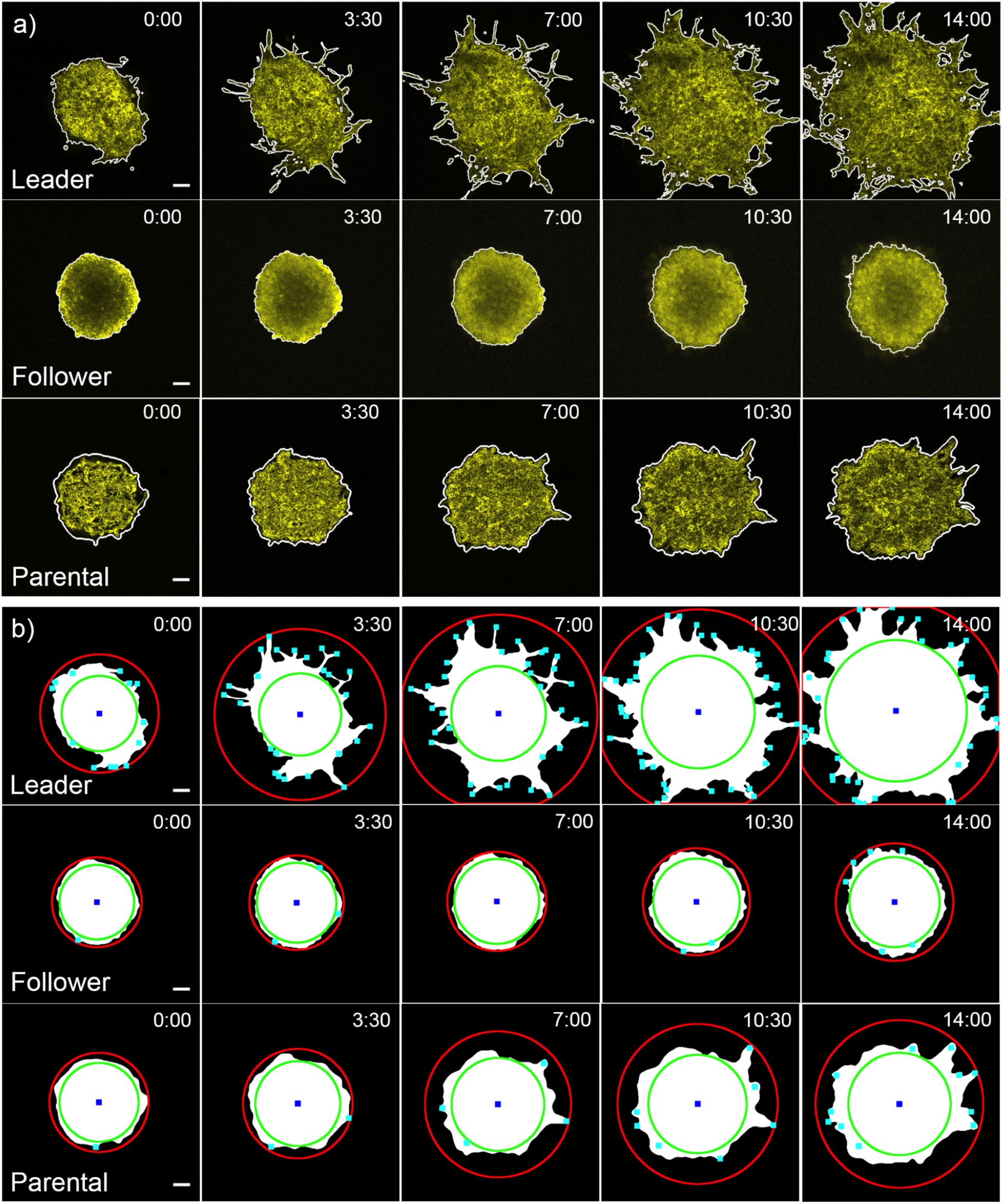
Distinct patterns of dynamic features. a) 3D graphcuts segmentation results. The white outlines are segmented boundaries for leader, follower and parental spheroids at different time points (Hour: Minute). Scale bars are 100 μm. b) Feature extraction results. The red circles are invasive radius; green circles are core radius; blue squares are centroids and cyan dots are branch points for leader, follower and parental spheroids at each time points (Hour: Minute). Scale bars are 100 μm.

### Preprocessing and image segmentation

Preprocessing methods are employed to enhance the contrast and salience of structures to improve image segmentation quality (see Figure 3). Four-dimensional volumes are first projected to 3D (*x, y, t*) using the standard deviation filter developed in [38]. At a given time t_0_, the volumetric image I(*x, y, z*_*i*_, *t*_*0*_) contains multiple focal planes *z*_*i*_ (see Figure 3.a). Uneven illumination of the focal plane in semi-solid gels can weaken the appearance of structures (see Figure 3.a, Z = 5, t_0_) making segmentation difficult. To correct this a standard deviation filter is applied at each time point to integrate focal planes and to define the 2D image sequence I_*p*_(*x, y, t*) = *σ*_*z*_ [I(*x, y, z, t*)]. These time-domain volumes are then Gaussian smoothed in both space (*x, y*) and time (*t*) to further mitigate noise. To delineate the spheroid boundaries, we leverage both spatial and temporal structure simultaneously by using an energy-minimizing graph cut segmentation [39-41]. The max-flow graph cut provides a smooth segmentation in both space and time using the similarities of spatial-temporal pixel neighbors. Treating these pixels as a spatial-temporal graph, where edge weight corresponds to inverse similarity, the algorithm finds an energy-minimizing cutting path through the volume to partition foreground and background regions. Example segmentations are provided in Figure 2.

**Figure 3.**
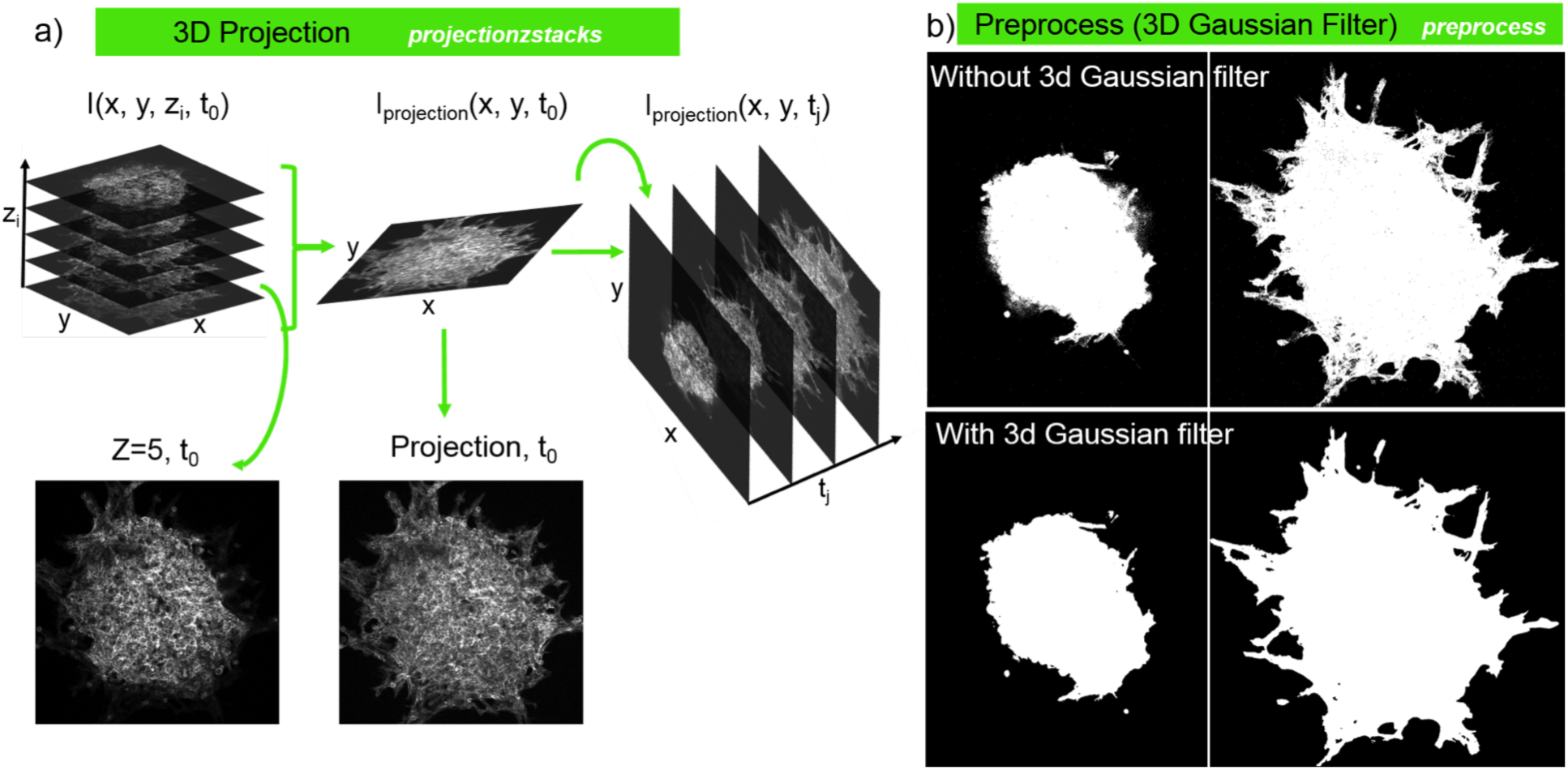
Contrast enhancement and smoothing. a) 3D projection algorithm converting z stacks at each time point to projection image, and then using projection images at each time point for preprocess. b) Segmentation effects comparison between without and with 3D Gaussian filter. 3D Gaussian filter smooth the segmentation in both spatial and temporal domains.

### Feature extraction

Given spheroid segmentation masks, TASI extracts a number of features to characterize static spheroid morphology at each time-point in the volume (see complete list, Table 2). Basic morphology features including area, perimeter, eccentricity, and intensity statistics are calculated. Complexity of the spheroid boundary is also measured as

**Table 2.** Statistical analysis of modeling parameters for spheroid types.

| Feature | Best model | ANOVA $p_a$<br>(linear) | ANOVA $p_b$<br>(quadratic) | Adjusted $R^2$ Range | | |
| --- | --- | --- | --- | --- | --- | --- |
|  |  |  |  | Leader | Follower | Parental |
| Area | Linear | **3.40e-7** | - | 0.993-0.995 | 0.990-1.0 | 0.997-1.0 |
| Invasive radius | Quadratic | **9.49e-3** | **3.90e-3** | 0.944-0.970 | 0.877-0.979 | 0.816-0.970 |
| Branch number | Quadratic | **1.12e-3** | **1.19e-4** | 0.594-0.867 | 0.553-0.904 | 0.840-0.902 |
| Core radius | Linear | **1.05e-3** | - | 0.909-0.951 | 0.945-0.982 | 0.921-0.992 |
| Perimeter | Quadratic | **7.96e-6** | **3.93e-9** | 0.877-0.980 | 0.930-0.976 | 0.936-0.989 |
| Complexity | Quadratic | **9.88e-8** | **7.16e-10** | 0.585-0.903 | 0.610-0.885 | 0.667-0.970 |
| Intensity mean | Quadratic | **7.15e-5** | **2.91e-3** | 0.990-0.990 | 0.980-0.993 | 0.990-0.998 |
| Intensity deviation | Quadratic | **8.35e-3** | **3.59e-2** | 0.709-0.981 | 0.756-0.883 | 0.885-0.986 |
| Eccentricity | Quadratic | 1.52e-1 | 7.02e-1 | 0.689-0.972 | 0.262-0.816 | 0.528-0.995 |
| Single cell count | Linear | 6.08e-2 | - | -0.012-0.413 | 0.010-0.951 | -1.28-0.786 |

**Bold:** $p < 0.05$

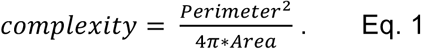

More irregular shapes will have larger perimeters for a corresponding area, translating to higher complexity measures (a circle has complexity 1).

Spheroids often exhibit interesting branching behavior, forming thin branches of invasive cells that protrude from the main spheroid mass. To quantify this phenomenon, we defined a “core radius” that captures the size of the main spheroid mass and an “invasive radius” that captures the extent of the projections (see Figure 1). The core radius was defined as the radius of the largest circle that can be inscribed within the spheroid mask, centered at the mask centroid. The invasive radius was defined from the minimum circle that can encompass the entire spheroid, including any invasive branches. These radii roughly capture growth due to proliferation and growth due to invasion. The number of the branches was further quantified using a skeletonization procedure. Morphological operations were applied to thin the mask to a skeletal structure, and the terminal endpoints were counted. This process robustly captures the tips of branching structures, even with complex shapes (see Figures 1, 2.b).

The presence of any isolated “cells” not connected from the main spheroid mass was also measured. These leaders are biologically extremely significant and may be represent a distinct cell phenotype with strong metastatic potential (see example Figure 1). These objects were detected by labeling the segmentation mask and looking for disconnected islands of foreground with a small area.

### Modeling and statistical analysis

The temporal evolution of measured features contains important information about spheroid dynamics and invasion. To measure dynamics, we provide mathematical modeling capabilities to fit models to temporal feature sequences. Three models are available for fitting: linear, quadratic, and exponential.

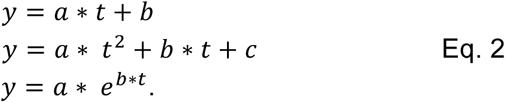

Modeling can be applied to individual spheroids, or to the average feature values of replicates for a single experimental condition. An adjusted R^2^ value is reported for each model as a measure of model fitness.

Statistical testing can be performed to on the model parameters to determine if there are measurable differences in spheroid dynamics across experimental conditions. Comparisons between pairs of treatments are made using the student’s t-test. Comparisons across two or more experimental conditions use the ANOVA test.

### Visualization and reporting

TASI automatically generates visualizations and reports for image analysis, model fitting, and statistical analysis results. For each experimental condition, time plots of the features are generated along with confidence intervals to illustrate variance within the condition replicates (see Figure 4). Feature plots and modeling can also be generated individually for each replicate in all experimental conditions (see Figure S2).

**Figure 4.**
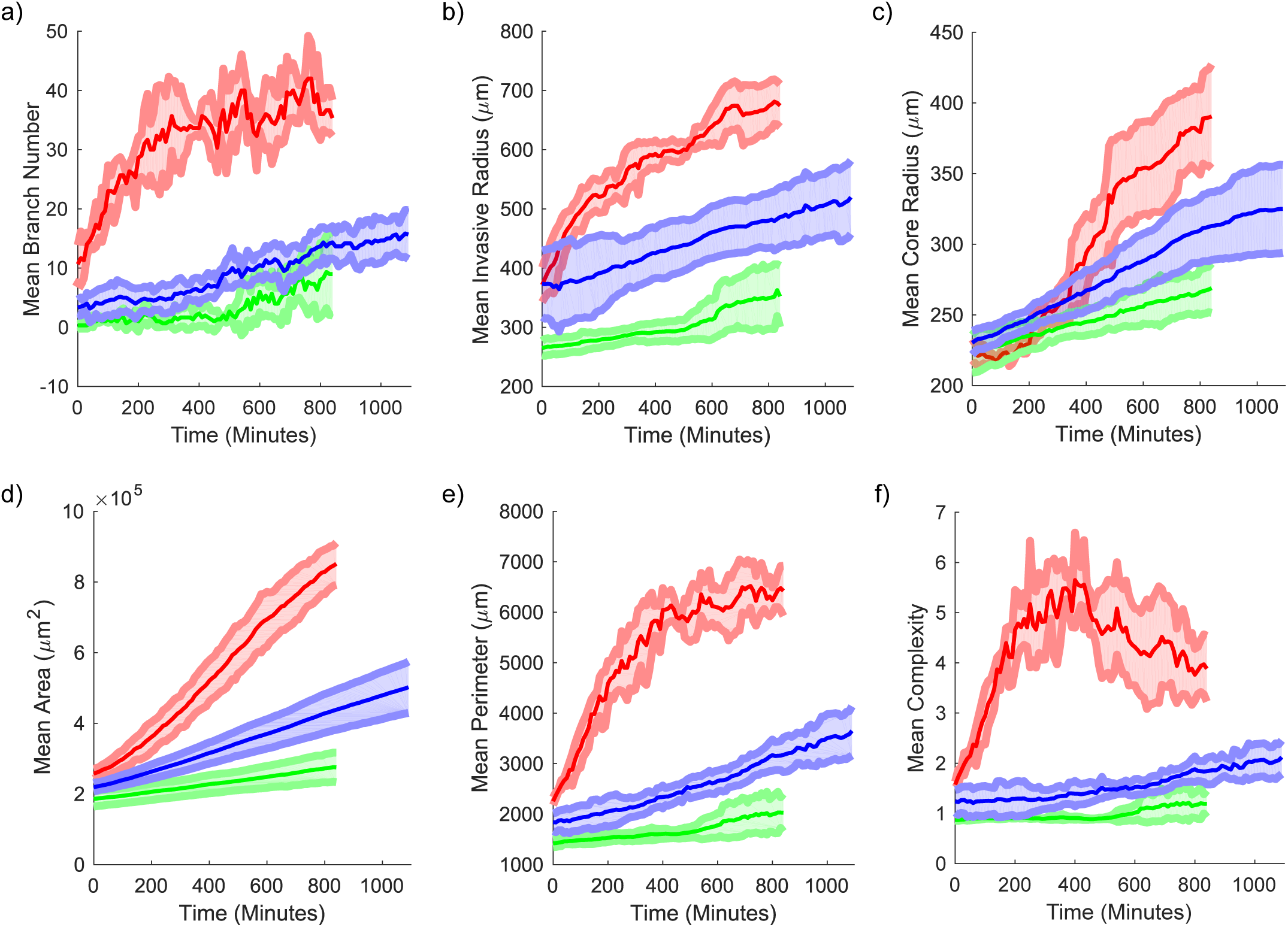
Visualization of average dynamic features under different treatments. Average values for a) branch number, b) invasive radius, c) core radius, d) area, e) perimeter and f) complexity as a function of time for different cell lines. Red solid lines with x symbols represent mean values for leader spheroids. Shaded areas represent 95% confidence intervals for feature values over the replicates.

## RESULTS

### TASI software

TASI is published as open-source software under an Apache 2.0 license (<u>https://github.com/cooperlab/TASI</u>). Full documentation on using TASI and the formatting of inputs and outputs is described in the Github repository. As an open-source framework, TASI can be readily and easily modified and extended to meet the needs of individual researchers.

As shown in Figure S1, TASI enables users to analyze individual spheroids, or spheroids grouped by experimental conditions. TASI uses a simple folder structure to organize and group experimental conditions, and performs end-to-end analysis once the input and output folders have been identified. Currently TASI supports most common image formats (jpg, png, tiff and others). The outputs generated for each spheroid include segmentation masks, feature extraction images, and spreadsheet CSV files containing feature extraction data and image analysis parameters for reproducibility. Additional CSV files describing modeling parameters and optionally statistical tests are also generated in the base output folder.

The spatio-temporal segmentation approach produces smooth segmentations of various spheroid morphologies (see Figure 2.a). By performing graph cuts simultaneously across both space (*x, y*) and time (*t*), the segmentation algorithm is able to suppress noise and produce segmentations that are smooth in both dimensions. Dim branches and edges that are characteristic of invasive spheroid phenotypes can be segmented accurately using this approach by integrating information across time. The graph-cutting approach can also compensate for gradual decreases in spheroid intensity over time due to imaging, as compared to segmentation methods that establish a uniform threshold for all time.

Segmentation masks are automatically saved in the output folders for quality control, where segmentation boundaries are superimposed over the spheroid intensity images in videos. By examining the video, the users can quickly review the segmentation quality, and refine if necessary.

### TASI analysis captures differences in spheroid phenotypes

Coordination between leader and follower cells in collective migration has been investigated, and key differences between leader and follower cells include cytoskeleton structure [42] and signaling pathway activations [29, 43, 44]. It is not clear, however, whether these complementary differences have a genetic basis or are induced by microenvironmental conditions. It is unknown, for example, if a follower cell becomes a leader when exposed to the spheroid/environment interface, or to what extent leader and follower phenotypes are transitory states that can be reversed. To answer these questions, we used TASI to analyze spheroids that were derived from purified leader or follower cells using <u>S</u>p<u>A</u>tiotemporal <u>G</u>enomic and cellular <u>A</u>nalysis (SaGA).

The morphological differences between these spheroids are apparent in Figure 2. Leader spheroids exhibit extensive chain-like branching, where the follower spheroids are compact and have more regular boundaries and sheet-like growth. The parental spheroids used to derive these populations are shown for comparison, and have intermediate morphologic qualities. Branch detection and core/invasive radius detection results for these spheroids are shown in Figure 2. b. The branch detection and radius finding algorithms work effectively across all spheroid types.

Temporal plots of key features are presented in Figure 4 for each spheroid type. We noted from the interval in these plots that trends are remarkably stable for each spheroid type, with replicates from a given type exhibiting little variation, suggesting that the segmentation and feature extraction methods are robust. Significant differences exist were observed in how the features evolved over time for each spheroid type. Leader spheroids rapidly develop branches in the first 5 hours of growth and then reach a plateau, although the core radius continues to increase. The branch number for follower spheroids increases very slowly and consistently over 8 hours, and proportional to core radius (and hence spheroid circumference). The complexity of follower spheroids remains close to 1 for all time points, suggesting a spherical morphology. Trends for parental spheroids were similar to follower spheroids, with a slight shift towards a more invasive phenotype.

### Model fitting and statistical testing of spheroid types

To further quantify differences in migration and growth between spheroid types, we fit models to the temporal sequences of each feature using least squares. Linear and exponential models are the most commonly used models for tumor growth, so we utilized these two models as well as a quadratic model to the sequences in Figure 5. Fitted models are shown in Figure 5. We noted that the R^2^ values are generally high, ranging from 0.71 to 0.99. Linear models accurately describing the temporal evolution of follower and parental spheroids. The temporal evolution of the leader spheroids is much more complex, and was better fit by the quadratic models in most cases (adjusted R^2^ ranges 0.71-0.99). The branch number and complexity dynamics of the leader spheroids are an exception, and are not described well by any model (adjusted R^2^ ranges 0.58 ∽ 0.90).

**Figure 5.**
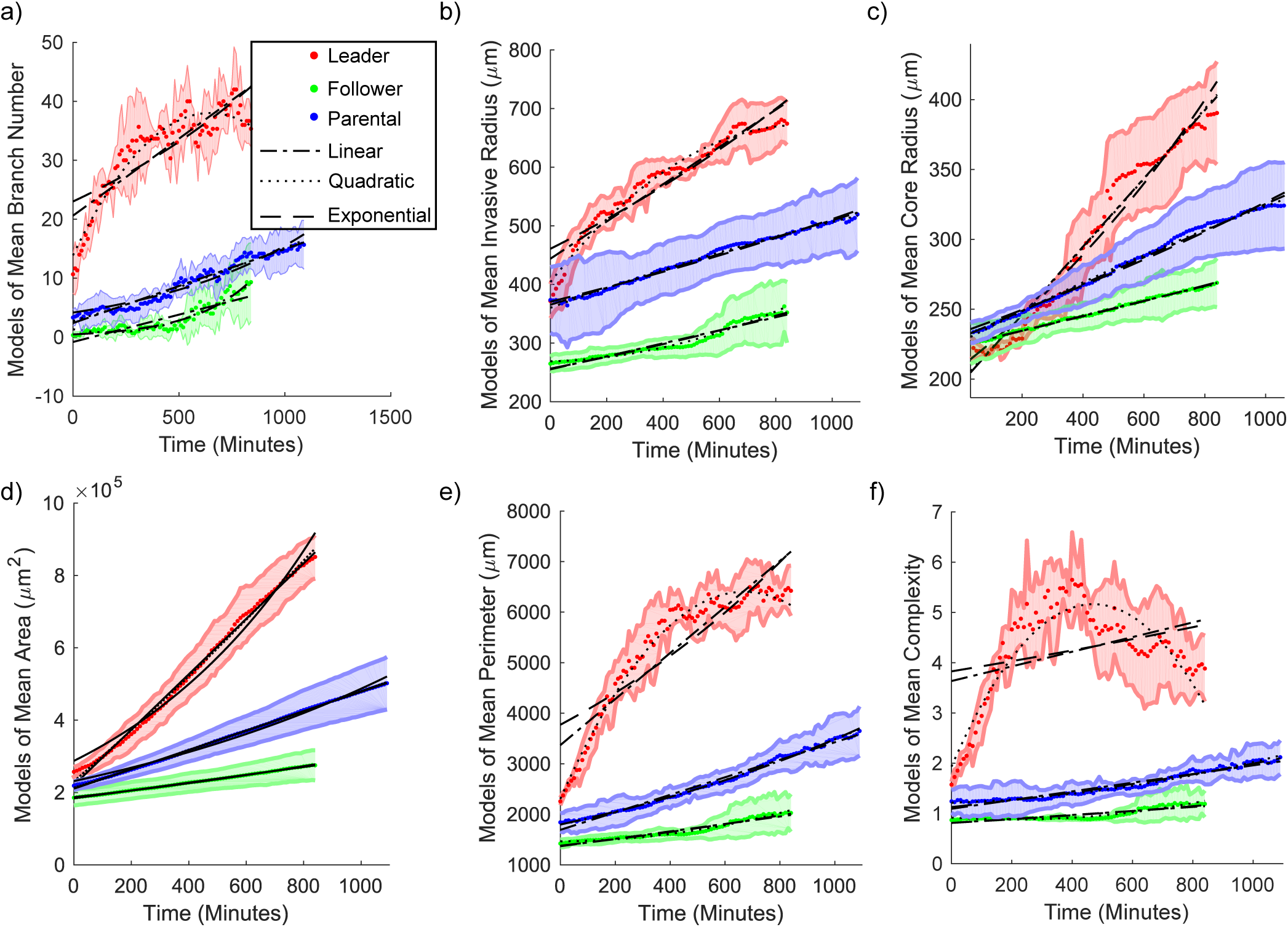
Model temporal feature evolution. Model fitting curves for mean a) branch number, b) invasive radius, c) core radius, d) area, e) perimeter and f) complexity as a function of time for each cell line. The dash-dot line represents the 1^st^ order polynomial (or linear) model fitting. The dotted line represents the 2^nd^ order polynomial model fitting. The dashed line represents the exponential model fitting. Shaded areas represent 95% confidence intervals for feature values over the replicates.

We performed statistical tests on these models to further quantify differences between the three spheroid types (see Table 2). The summary of the statistical test results among three cell lines is listed in Table 2. The features with the most significant differences in model parameters included area (linear model, ANOVA p = 3.40e-7), complexity (quadratic model, p_a_ = 9.88e-8, p_b_ = 7,16e-10), and perimeter (quadratic model, p_a_ = 7.96e-6, p_b_ = 3.93e-9). Eccentricity and single cell number had the weakest differences across spheroid types. The statistical tests confirmed our aim of defining unique features to classify different collective migration patterns.

## DISCUSSION

The application of TASI image analysis to spheroid data obtained by SaGA illustrates how spheroid cultures and image analysis can be used to investigate tissue microenvironments and their role in cancer invasion and metastasis. TASI contributes an end-to-end software approach for characterizing spheroid growth and invasion dynamics, providing image segmentation, feature extraction, modeling, and statistical analysis capabilities within the same tool. As an open source framework, it can readily be extended and tailored to the specific needs of investigators.

Our analysis of spheroids derived from purified leader and follower cells [37] used features like branch tip count and core radius to reveal important differences in growth and invasion. Leader and follower cells likely play complementary roles in establishing viable metastases distant from the primary tumor, and understanding this process and the differences in these cell phenotypes can lead to better targeting of these important mechanisms in the future. Growth and invasion are by definition dynamic processes, and by providing a framework to measure dynamic behaviors of spheroids, TASI enables precise and quantitative characterization of spheroid behavior. The ability to model these behaviors and to perform statistical tests between experimental conditions could aid in screening drugs or functional genetics studies by detecting subtle differences in dynamic behavior as opposed to static morphology. Objectivity and repeatability in these types of experiments is increasingly critical, and with large amounts of data, traditional manual quantification may not be practical.

In the future we plan to extend TASI to perform true three-dimensional temporal imaging (x,y,z + time) of spheroid cultures, although we see from our experiments that differences in spheroids were clearly measurable despite projecting/flattening (x,y,z) volumes. Extending TASI to leverage existing cell tracking algorithms is another important direction for future development. Tracking will enable more detailed analysis of the movement patterns of leader cells, and more complex characterizations of the constituents in chain-like projections.

## ACKNOWLEDGEMENTS

This work was supported by U.S. National Institutes of Health, National Library of Medicine Career Development Award K22LM011576, and National Cancer Institute grants U24CA180924 and U24CA194362.

## AUTHOR CONTRIBUTIONS

Y.H. and L.C. developed the software framework and algorithms and performed documentation and installation packaging. J.K. performed experiments and imaging. Y.H. performed image analysis of experimental data. D.B., A.I.M. and L.C. assisted with manuscript development. L.C. conceived of the software concept and supervised the project.

